# Focal Transcranial Magnetic Stimulation of the Rat Anterior Cingulate Cortex Inhibits Incubation of Opioid Craving after Voluntary Abstinence

**DOI:** 10.64898/2026.03.04.709400

**Authors:** Zilu Ma, Ying Duan, Hieu Nguyen, Sabrina Lin, Md Mohaiminul Haque, Danni Wang, Samantha Hoffman, Aidan Carney, Taylor Scott, Olivea Varlas, Elliot A. Stein, Zheng-Xiong Xi, Yavin Shaham, Hanbing Lu, Yihong Yang

## Abstract

Relapse remains a major challenge in opioid addiction treatment, underscoring the need for innovative therapies. Progress in neuromodulation therapies has been limited by insufficient mechanistic understanding of stimulation engagement and disease-related changes in the brain.

We used a novel, focal transcranial magnetic stimulation (TMS) system to deliver high-density theta burst stimulation (hdTBS) combined with resting-state fMRI to test whether anterior cingulate cortex (ACC) stimulation reduces relapse-like behavior and alters functional circuitry in a rodent model of opiate dependence. The coil focality and stimulation parameters approximate human TMS protocols, and the targeted region represents a functional homolog of the human ACC.

We trained rats to self-administer oxycodone intravenously for 14 days. We then introduced an electric barrier for 13 days, which caused cessation of drug self-administration. We assessed relapse to oxycodone seeking immediately after training (early abstinence) and after electric-barrier exposure (late abstinence). We administered daily hdTBS or sham stimulation for 7 days before the late-abstinence test.

Sham-treated rats showed a time-dependent increase in oxycodone seeking during abstinence (incubation of oxycodone craving) and reduced ACC functional connectivity. In contrast, hdTBS prevented the incubation of oxycodone craving and restored ACC connectivity with the dorsal and ventral striatum. Tracer-based axonal-projection data further showed that stimulation-induced effects aligned with regions receiving dense projections from the stimulation site, suggesting that the projection architecture is critical to the propagation of focal stimulation across distributed networks.

These findings identify ACC-centered circuits as mechanistically informed targets for TMS-based interventions that aim to reduce opioid relapse during abstinence.

**One sentence Summary:** Prefrontal TMS stimulation reduced relapse-like behavior and restored corticostriatal circuits, highlighting translational targets for addiction treatment.

## INTRODUCTION

Drug addiction is a major global health challenge affecting more than 60 million people worldwide (UNODC, 2025). In the United States, opioid-related overdoses account for ∼76% of all drug-related deaths, representing a significant public health burden (Garnett & Minino, 2024). A defining feature of addiction is relapse to drug use after treatment cessation (Jaffe, 1990). Relapse risk persists after prolonged abstinence and remains a major treatment challenge (Hunt et al., 1971; Sinha et al., 2011). In humans, re-exposure to drug-associated cues and contexts often triggers craving and relapse (Garavan et al., 2000; O’Brien et al., 1992). In rodents, relapse to opioid seeking increases with abstinence duration, a phenomenon termed *incubation of drug craving* (Chow et al., 2025; Grimm et al., 2001). Incubation occurs after both forced homecage abstinence (Grimm et al., 2001; Pickens et al., 2011; Shalev et al., 2001) and voluntary abstinence induced by exposure to an electric barrier (Fredriksson et al., 2020; Fredriksson, Tsai, et al., 2021; Negishi et al., 2024).

Extensive research has focused on pharmacological and behavioral treatments for opiate addiction (Carroll & Onken, 2005; Venniro et al., 2020; Volkow et al., 2016). Three FDA-approved medications targeting the μ-opioid receptor, methadone, buprenorphine, and naltrexone, are currently available (Cunningham et al., 2020; Skolnick, 2018). Although these treatments reduce opioid use, relapse, and overdose, relapse rates remain high due to limited treatment access and individual variability in treatment response (Epstein et al., 2018; Kampman & Jarvis, 2015; Weiss et al., 2011). In animal models, drugs targeting dopamine, glutamate, noradrenaline, and other neurotransmitter systems reduce, at least partially, some relapse-related behaviors (Bossert et al., 2013; Chow et al., 2025), but these findings have not yet translated into efficacious FDA-approved therapies (Venniro et al., 2020). This translational gap underscores the need for treatments that reverse drug-induced neuroadaptations hypothesized to be critical to relapse vulnerability (Kalivas et al., 2009; Wolf, 2025).

Brain stimulation has emerged as a potential strategy to reverse addiction-related neuroadaptations (Ekhtiari et al., 2019). However, clinical trials using deep brain stimulation, transcranial direct current stimulation, and transcranial magnetic stimulation (TMS) have largely relied on trial-and-error approaches and have produced mixed results (Dinur-Klein et al., 2014; Hanlon et al., 2016; Luigjes et al., 2019; Steele & Maxwell, 2021), likely due to clinical heterogeneity, including polysubstance use, psychiatric comorbidity, and concurrent medications, which complicate interpretation of stimulation outcomes (Volkow et al., 2016). Preclinical addiction models offer key advantages, including control over drug exposure, genetics, and environment, and facilitate longitudinal assessment across addiction phases (Creed, 2018). These models also permit invasive and non-invasive neuromodulation with circuit-level precision, enabling causal tests of treatment mechanisms (Spanagel, 2017).

While preclinical brain stimulation studies are critical for optimizing clinical interventions, applying TMS to small animals involves multiple technical challenges (Wilson et al., 2018). A small coil is necessary to reduce off-target effects (Deng et al., 2013). However, such coils are limited by low efficiency, overheating, and excessive electromagnetic stress (Cohen et al., 1990). Thus, previous preclinical studies have used oversized coils that likely stimulated most of the brain or miniature coils that were too weak to elicit action potentials (Alekseichuk et al., 2019; Gersner et al., 2011; Parthoens et al., 2016; Rotenberg et al., 2010; Tang et al., 2016; Vahabzadeh-Hagh et al., 2012).

Our group recently developed and extensively validated a novel coil design that overcomes these limitations, enabling highly focal (< 2 mm), suprathreshold stimulation of the rodent brain, which allows selective modulation of targeted brain regions at a human-comparable scale (Cermak et al., 2020; Meng et al., 2018; Meng et al., 2022). We also developed a high-density theta-burst stimulation (hdTBS) technology that delivers six pulses per burst, doubling the conventional TBS pattern (Huang et al., 2005) and enhancing acute after-effects by approximately 92% (Meng et al., 2022).

In the current study, we used this focal TMS coil and hdTBS to examine the effect of stimulating the anterior cingulate cortex (ACC) on the incubation of opioid craving after voluntary abstinence in rats. The ACC is a central hub within the frontostriatal network circuits, interconnected extensively with striatum and insula (Haber & Knutson, 2010; Sutherland & Stein, 2018), plays a critical role in reward valuation, action monitoring and cognitive control (Rushworth & Behrens, 2008; Shenhav et al., 2013). Dysregulation of ACC to striatal circuitry has been implicated in addiction and relapse vulnerability (Li et al., 2017; Sutherland & Stein, 2018). In a recent study, we combined functional MRI (fMRI) with pharmacological inactivation and identified a critical role of frontostriatal circuitry in relapse to oxycodone seeking after voluntary abstinence (Ma et al., 2025). Specifically, inactivation of the dorsomedial striatum (DMS) reduced oxycodone seeking after electric barrier-induced abstinence, and this effect was associated with increased functional connectivity between the ACC and DMS. Based on these findings, as well as earlier studies on the role of ACC and dorsomedial PFC in cue- and context-induced relapse to opioid seeking in rat models (McKendrick et al., 2022; Rogers et al., 2008), we used the ACC as a stimulation target to investigate whether TMS stimulation during voluntary abstinence would decrease incubation to oxycodone seeking.

We trained rats to self-administer oxycodone for 14 days and then subjected them to 13 days of electric barrier-induced voluntary abstinence. This approach models abstinence by imposing an electric barrier of increasing intensity in front of the drug-paired lever while maintaining intact response-outcome contingencies, thereby approximating adverse consequence-driven, clinically relevant patterns of drug cessation (Fredriksson et al., 2020; Fredriksson, Tsai, et al., 2021). We tested relapse to oxycodone seeking on abstinence day 1 (early), immediately after self-administration training, and on abstinence day 15 (late), after 13 days of voluntary abstinence. We administered daily hdTBS or sham stimulation for seven days before the late-abstinence relapse (incubation) test. To identify circuit-level mechanisms, we conducted longitudinal functional MRI (fMRI) at early and late abstinence (Fredriksson, Tsai, et al., 2021) and compared abstinence-related changes in functional connectivity between the hdTBS and sham groups.

## RESULTS

### hdTBS stimulation of ACC decreased incubation of oxycodone seeking after electric barrier-induced voluntary abstinence

We trained rats (n = 36) to self-administer oxycodone (0.1 mg/kg) under fixed-ratio 1 (FR1) schedule for 14 days (6 h/day), followed by 13 days of electric barrier-induced voluntary abstinence (2 h/day; Figure 1A). All rats demonstrated reliable oxycodone self-administration, as indicated by a progressive increase in the number of infusions earned across training sessions (Figure 1B left panel, Table S1, Training Session, F_13, 442_ = 89.62, p < 0.001). During the electric-barrier phase, the rats voluntarily abstained from oxycodone self-administration as shock intensity was progressively increased from 0 mA to 0.4 mA (Figure 1B right panel, Table S1, Electric Barrier Session, F_12, 408_ = 104.25, p < 0.001).

**Figure 1.**
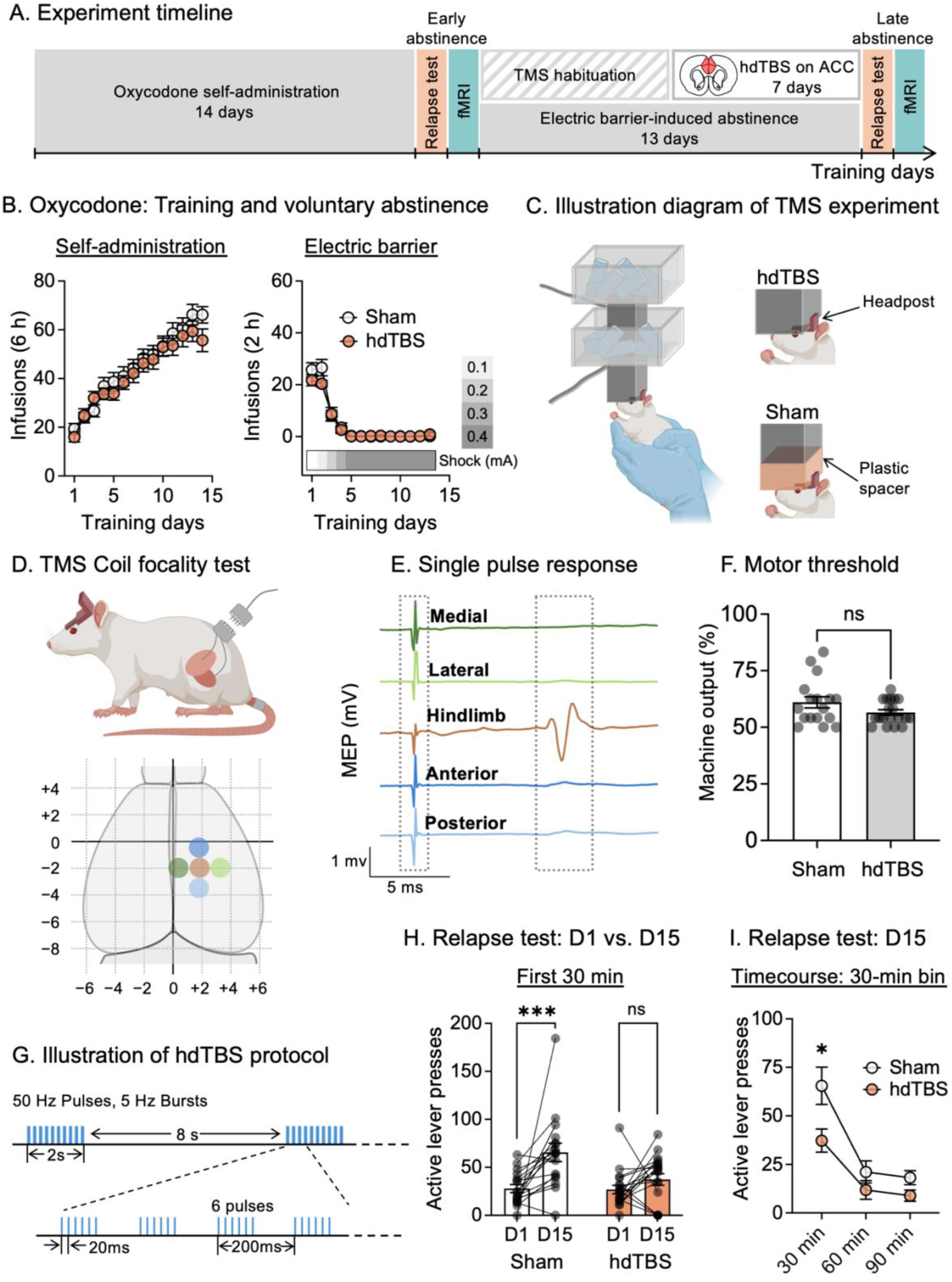
Chronic TMS stimulation reduces incubation of oxycodone seeking. (**A**) Experimental timeline.(**B**) Oxycodone self-administration training (left) and electric barrier-induced voluntary abstinence (right). Data show mean ± SEM of number of infusions for sham and hdTBS rats. (**C**) Schematic diagram of TMS stimulation. Created with Biorender.com. (**D-E**) Coil focality mapping based on hindlimb motor-evoked potentials (MEPs) during single pulse stimulation with 1 mm offsets. Dashed boxes indicate stimulus artifact window (left) and MEP response window (right). Data replotted from (Meng et al., 2022). (**F**) Motor threshold for sham and hdTBS rats. (**G**) Illustration of hdTBS stimulation protocol. Six pulses per burst delivered at 50 Hz (20 ms interpulse interval). Ten bursts per train at 5 Hz (200 ms interburst interval). Twenty trains (2 s each) per session separated by 8 s intertrain intervals (1200 total pulses over 192 s per session). (**H**) Relapse test: day 1 vs. day 15. Data show mean ± SEM number of active lever presses during the 30-min test of day 1 and the first 30-min of the test on day 15. (**I**) Relapse test day 15: 30 min bin timecourse. ***p < 0.001, *p < 0.05. n = 17 sham, n = 19 hdTBS. fMRI: functional MRI; hdTBS: high-density theta burst stimulation; ACC: anterior cingulate cortex; MEP: motor-evoked potential; D1: day 1; D15: day 15.

Using a custom-built TMS coil and the hdTBS stimulator (Figure 1C) (Cermak et al., 2020; Meng et al., 2018; Meng et al., 2022), we assessed the focality of TMS delivery by measuring the motor-evoked potentials (MEPs) during single-pulsive stimulation while offsetting the coil position by approximately 1 mm along four directions (medial, lateral, anterior, and posterior) around the hindlimb region of the motor cortex (Figure 1D), as previously described (Meng et al., 2022). As shown in Figure 1E, the MEP amplitudes were maximal when the coil was centered over the hindlimb motor cortex and decreased substantially with 1 mm offsets.

On abstinence day 1, we tested the rats for oxycodone seeking under extinction conditions for 30 min (Figure 1A). During testing, pressing the active lever activated the tone and light cue previously associated with oxycodone infusions but no drug infusions were delivered. On abstinence days 2-7, we habituated the rats to the TMS procedure (three times per day, at least 1 h before electric barrier training) to minimize motion and stress during subsequent stimulation (Figure 1C, see Materials and Methods for additional details). The rats remained awake, and were gently restrained by hand during stimulation (Figure 1C).

Following motor threshold determination, we randomly divided the rats into sham (n = 17) and active hdTBS (n = 19) groups (see Materials and Methods for additional details). No significant difference in motor thresholds was found between the sham and the hdTBS groups (Figure 1F, p = 0.10, t _34_ = 1.69). During abstinence day 8 to 14, we delivered either sham or TMS stimulation, using the hdTBS protocol (Figure 1G, see Materials and Methods for additional details), to the ACC one session per day (1200 pulses over 192 s) for 7 consecutive days, at least 1 h before the electric barrier sessions. The rats in both the sham and hdTBS groups showed a normal increase in body weight (Figure S1, Table S1), with no changes in eating, drinking, grooming, or locomotor behavior. In addition, we observed no TMS-induced motor seizures during or after TMS administration. On abstinence day 15, we tested the rats for relapse to oxycodone seeking under extinction conditions for 90 minutes.

Sham-treated rats showed a time-dependent increase in active lever presses from day 1 to day 15 (incubation of oxycodone seeking), and this effect was absent in the active ACC hdTBS group (Figure 1H). We analyzed active lever presses during the 30-min relapse test on day 1 and the first 30-min of the relapse test on day 15 with RM-ANOVA using the between-subjects factor of Group (sham, hdTBS) and the within-subject factor of Abstinence Day (day 1, day 15). This analysis showed a significant Group × Abstinence Day interaction (Figure 1H, F_1, 34_ = 4.40, p = 0.043). Additionally, we analyzed the full 90-min test data from day 15 with RM-ANOVA using the between-subject factor of Group and the within-subject factor of Session Time (30, 60, 90 min). This analysis showed a significant effect of Group (Figure 1I, F_1, 34_ = 7.36, p = 0.010) and Session Time (F_2, 68_ = 35.83, p < 0.0001) but no significant Group × Session Time interaction (F_2, 68_ = 2.42, p = 0.097, Table S1), with the active hdTBS group showing significantly lower active lever presses during the first 30 min of the test session (Table S1).

We performed the same analysis on inactive lever presses (a measure of non-specific activity) during the 30-min test on day 1 and the first 30-min test on day 15. In contrast, we found no significant Group × Abstinence Day interaction (Figure S2, F_1, 34_ = 0.21, p = 0.65). Similarly, inactive lever presses in the 30-min bin timecourse analysis only showed significant effect of Session Time (Figure S2, F_2, 68_ = 13.72, p < 0.001) but no effect of Group (F_1, 34_ = 0.59, p = 0.45) nor Group × Session Time interaction (F_2, 68_ = 0.32, p = 0.73, Table S1).

Together, these results indicate that the active hdTBS prevented the emergence of incubation of oxycodone seeking after electric barrier-induced abstinence. The observation that active lever responding in the hdTBS group on day 15 was not different than day 1 suggests that the TMS effect is not due to non-selective deficits in operant responding but rather reflects a specific effect on ‘incubated’ oxycodone seeking.

### hdTBS stimulation of ACC reversed functional connectivity changes after electric barrier-induced voluntary abstinence

Prior human and rodent studies demonstrated that reduced functional coupling between the ACC and striatum and other limbic regions is associated with opioid addiction and relapse (Gorka et al., 2014; Ma et al., 2025; Upadhyay et al., 2010). Based on this evidence, we next used fMRI to examine the functional connectivity changes within the ACC network during voluntary abstinence (early to late abstinence) and the effect of hdTBS on these changes. We collected resting-state fMRI scans on the days following the early and late abstinence relapse tests (Figure 1A, day 2 and day 16). A total of 35 rats (n = 17 sham, n = 18 hdTBS) successfully completed fMRI data collection and passed data quality control (see Methods and Materials for more details).

We conducted a voxel-wise functional connectivity analysis using ACC as the seed region (Figure 2A left panel, AP: +3.5 ∼ +1.5 mm), followed by linear mixed-effect modeling with two factors: Group (sham, hdTBS) and Abstinence Day (day 2, day 16) to assess the differences in the time-dependent changes between the sham and hdTBS groups during the voluntary abstinence period. As shown in Figure 2A, several brain regions, including the dorsal striatum (DS), nucleus accumbens (NAc), motor cortex (Mc), and sensory cortex (Sc), showed a significant Group × Abstinence Day interaction effect in functional connectivity with the ACC (p < 0.05, corrected for multiple comparisons). We extracted functional connectivity values from these regions showing the significant Group × Abstinence Day interaction and plotted these values for individual rats in Figure 2B. In the ACC-DS and ACC-NAc circuits, the interaction effect was driven by a decrease in functional connectivity across abstinence in the sham group, whereas the active hdTBS decreased or reversed this pattern. In contrast, for ACC-Mc and ACC-Sc circuits, the interaction was reduced across abstinence in the active hdTBS group, with no change observed in the sham group.

**Figure 2.**
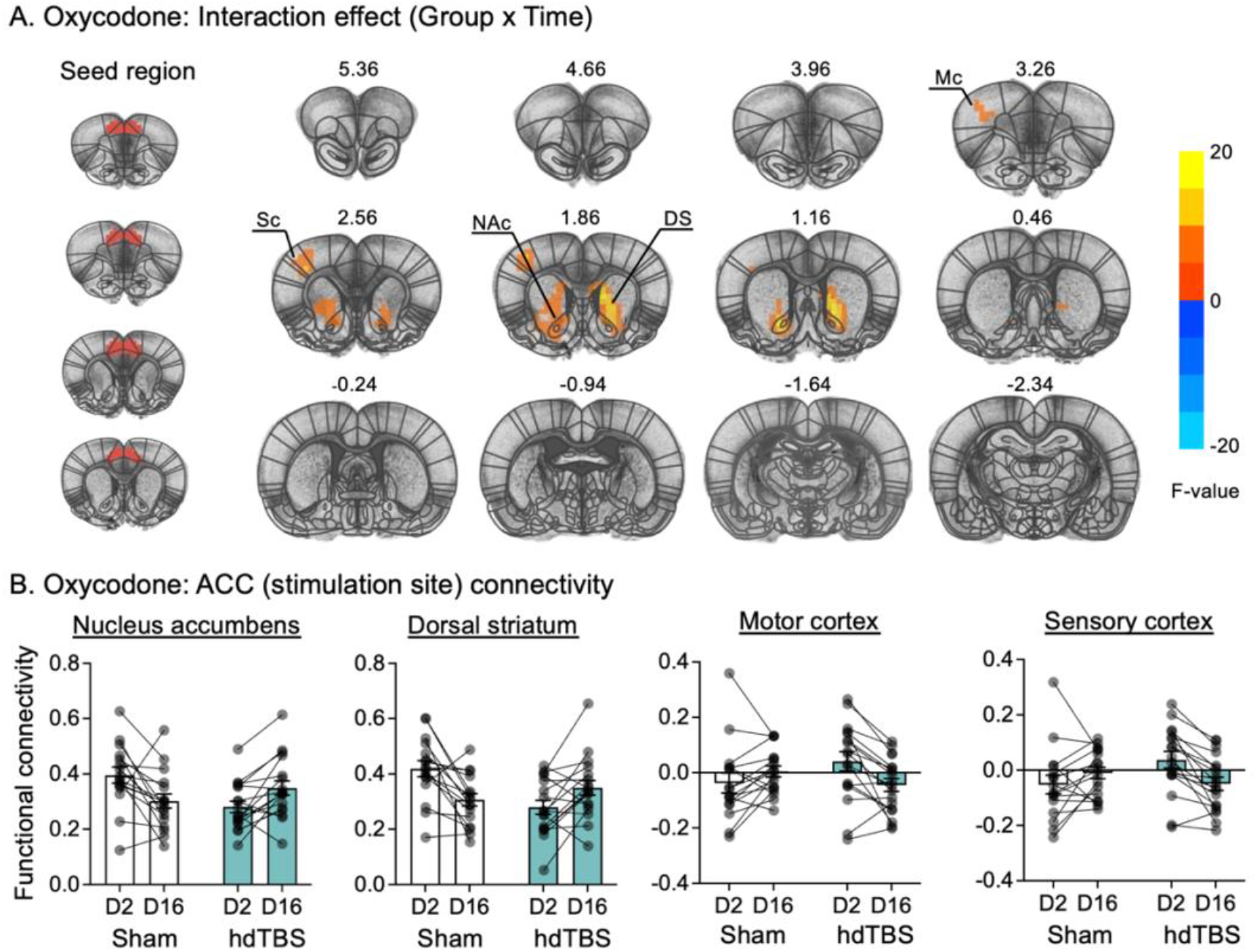
Effect of hdTBS to the ACC on brain functional connectivity. (**A**) Left panel: ACC seed region. Right panel: Group (sham, hdTBS) by Abstinence Time (early, late-abstinence) interaction in whole brain voxel-wise ACC (AP: + 3.5 ∼ +1.5 mm) functional connectivity (corrected p < 0.05, uncorrected p < 0.01, cluster size > 25). (**B**) Functional connectivity of individual rats between ACC and four brain regions from **A**: nucleus accumbens, dorsal striatum, motor cortex, and sensory cortex. Color-scale indicates F values. Corrected for multiple comparison. Data show mean ± SEM. n = 17 sham, n = 18 hdTBS. Mc: motor cortex; Sc: sensory cortex; NAc: nucleus accumbens; DS: dorsal striatum; ACC: anterior cingulate cortex; hdTBS: high-density theta burst stimulation; D2: day 2; D16: day 16.

To further examine these changes, we next performed a post-hoc analysis on the difference in the ACC functional connectivity between early (day 2) and late (day 16) abstinence for each group. In sham rats, voxel-wise t-test showed a significant reduction in ACC functional connectivity with NAc and DS during the electric barrier-induced voluntary abstinence (Figure 3A, p < 0.05, corrected for multiple comparisons). In contrast, this reduction was not observed in the hdTBS treated group. Instead, there was a slight reduction in ACC connectivity with sensory and motor cortex in the hdTBS-treated group (Figure 3B, p < 0.05, corrected for multiple comparisons). Extracted functional connectivity values from these regions showing significant contrast were plotted for individual rats in Figure 3C and 3D, illustrating the direction and magnitude of observed effects.

**Figure 3.**
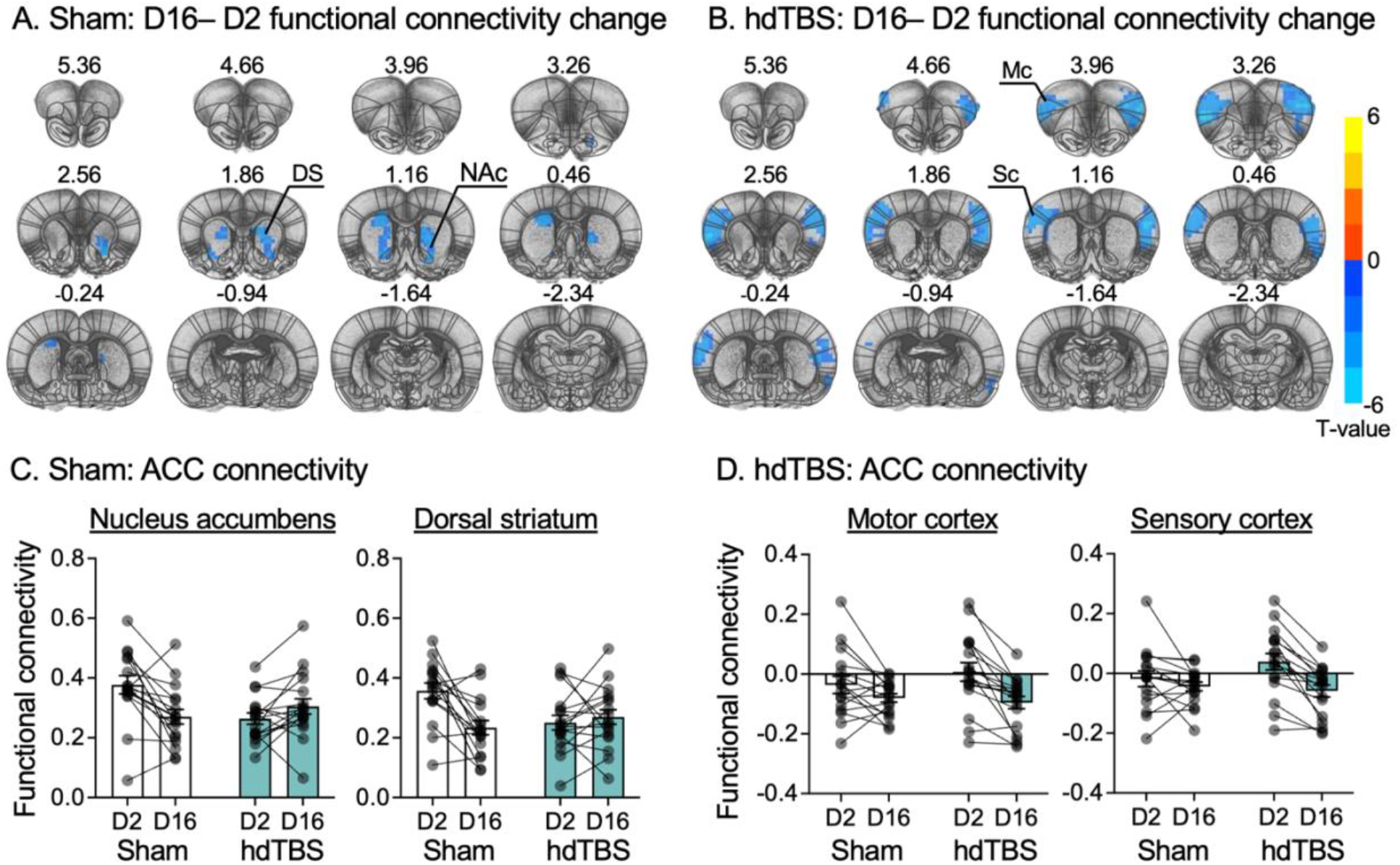
Post-hoc analysis on the effect of hdTBS stimulation on functional connectivity. (**A & B**) Post-hoc t-test following Group by Abstinence Time interaction in whole brain voxel-wise functional connectivity between early and late abstinence in sham (**A**) and hdTBS (**B**) group. (**C & D**) Functional connectivity of individual rats between ACC and brain regions extracted from voxelwise t-test in **A&B**: **C** nucleus accumbens and dorsal striatum; **D** motor cortex and sensory cortex. Color-scale indicates Z values. Corrected for multiple comparisons. Data show mean ± SEM. Sham n = 17, active n = 18. D2: day 2; D16: day 16; DS: dorsal striatum; NAc: nucleus accumbens; Mc: motor cortex; Sc: sensory cortex; ACC: anterior cingulate cortex; hdTBS: high-density theta burst stimulation.

These results indicate that electric barrier-induced abstinence decreased ACC-striatum functional connectivity, and this effect was prevented by repeated hdTBS stimulation of the ACC.

### TMS-induced network reorganization follows anatomical projection

To examine whether the TMS-induced circuitry changes align with known efferent pathways of the stimulated region, we compared the spatial pattern of the Group (sham vs. hdTBS) by Abstinence Day (early-vs. late-abstinence) interaction map with axonal projection maps from tracer injections into the ACC. Axonal projection data were adopted from the Allen Mouse Brain Connectivity Atlas (Oh et al., 2014) (https://connectivity.brain-map.org; Experiment ID: 112458114).

We first compared the spatial distribution of the ACC axonal projection with TMS-induced differential changes in functional connectivity (Group × Abstinence Day interaction; Figure 4A). These changes in functional connectivity correspond to the known spatial distribution of dense axonal projection sites from the ACC tracer-derived projection map, including the DS, NAc, and sensory/motor cortex (SM; Figure 4A). For quantitative comparison, we extracted the projection volume, projection density, as well as the F-values from the Group by Abstinence Day interaction of functional connectivity for regions including the prefrontal cortex (PFC), retrosplenial cortex (RSP), sensory/motor cortex (SM), lateral cortical area (CTX), DS, NAc, thalamus (TH), and hypothalamus (HY). Consistent with the spatial maps, the fMRI and the projection density matrix showed a high congruency (Figure 4B). We next assessed the percentage of projected volumes for each of these regions among all ACC projected volumes. We found that the combined projection volume of hdTBS-altered regions, DS, NAc, and SM, was greater than 55% of total ACC projected volume (Figure 4C).

**Figure 4.**
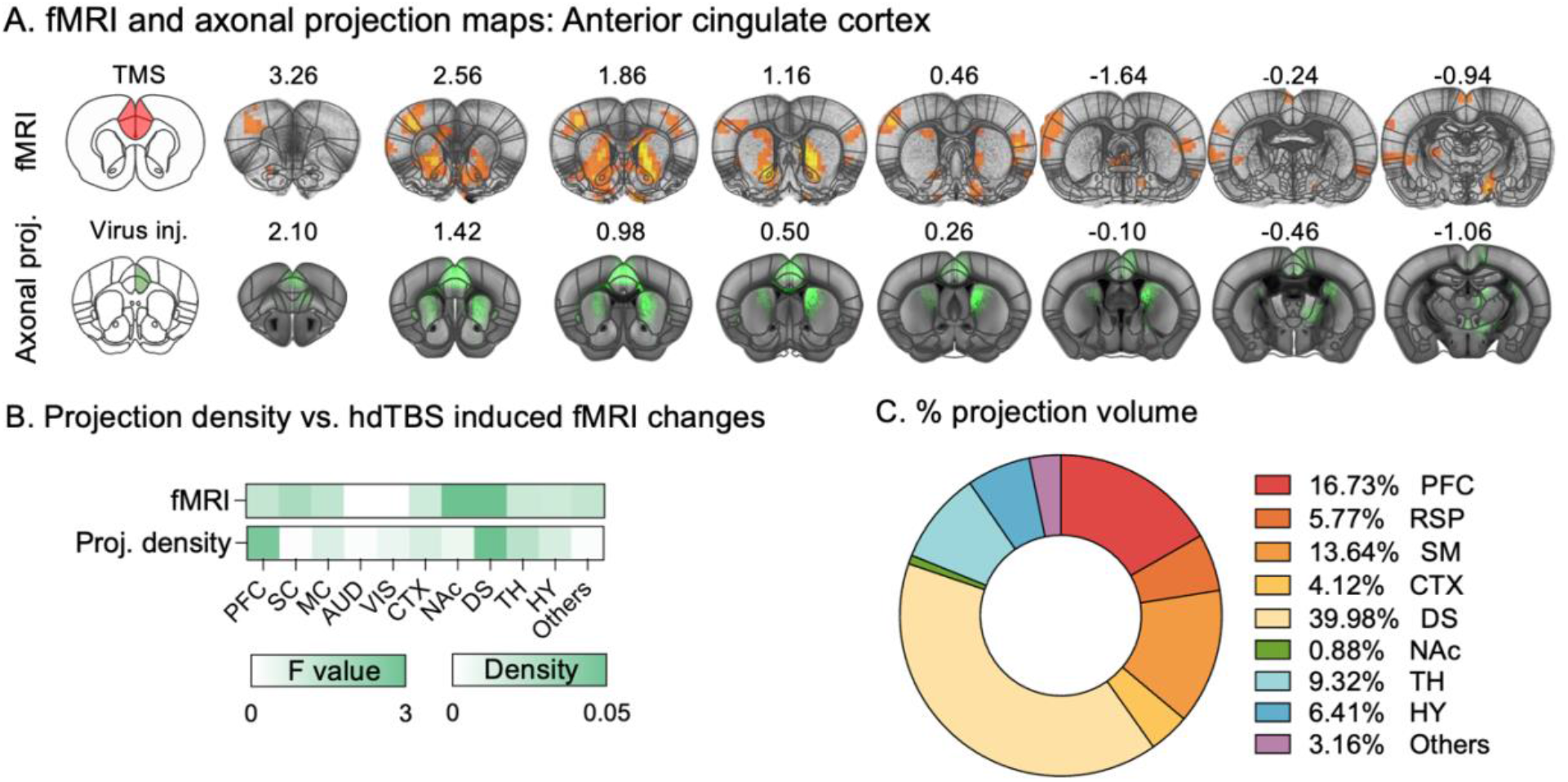
Tracer-based axonal projection vs. TMS-induced functional connectivity changes. (**A**) Top: Group by time interaction in whole brain voxel-wise functional connectivity of ACC (uncorrected p < 0.05). Color-scale indicates F values. Bottom: Tracer-based axonal projection map following AAV-EGFP viral injection into ACC (Experiment ID: 112458114). Axonal projection data were adopted from the Allen Mouse Brain Connectivity Atlas (Oh et al., 2014) (https://connectivity.brain-map.org). (**B**) Projection density and F values from Group by Abstinence Day interaction analyses of functional connectivity from ACC. (**C**) Percentage of regional projection volume over all projected volumes from ACC. PFC: prefrontal cortex; RSP: Retrosplenial cortex; SM: sensory/motor cortex; CTX: lateral cortical region; DS: dorsal striatum; TH: thalamus; HY: hypothalamus; Others: other brain regions.

Together, these results suggest that the efficacy and spatial specificity of ACC hdTBS may, at least, partially be determined by the underlying axonal projection architecture of the targeted regions, providing a mechanistic link between stimulation site selection and downstream circuit engagement.

## DISCUSSION

We tested the effects of hdTBS delivered with a custom-developed focal TMS coil on the incubation of oxycodone seeking after electric barrier-induced abstinence. We found that hdTBS directed to the ACC suppressed the incubation of oxycodone craving and prevented the reduction in ACC functional connectivity with dorsal striatum and nucleus accumbens from early to late abstinence. By comparing TMS-induced functional connectivity changes with tracer-based axonal projection maps from ACC, we demonstrated that regions exhibiting significant stimulation-dependent interaction effects preferentially overlapped with regions receiving dense axonal projections from the stimulation site. Importantly, the hdTBS treatment was well tolerated by the rats, with no behaviorally detected seizures or adverse side effects, supporting the safety and feasibility of focal neuromodulation in this model.

### ACC-centered circuitry as a translational target for neuromodulation-based intervention of drug craving

Comparative anatomical studies indicate that the rodent ACC shares similarities with the human dorsal, ventral, and rostral ACC (Brodmann areas 24, 25, and 32), regions implicated in decision-making, emotion regulation, conflict monitoring, and attentional control (van Heukelum et al., 2020; van Hout et al., 2024; Vogt & Paxinos, 2014). In people with drug addiction, these ACC subdivisions show structural and functional changes, including reduced gray matter volume, hypoactivation, and heightened cue-induced activation (Camchong et al., 2013; Goldstein et al., 2009; Goldstein & Volkow, 2011; Hu et al., 2015; Li et al., 2017; Wang et al., 2012). At the circuit level, the ACC show direct projections to the dorsal and ventral striatal regions (Haber & Knutson, 2010), and resting-state functional connectivity analyses showed intrinsic functional coupling between these regions in humans (Di Martino et al., 2008).

These ACC-striatal pathways therefore provide a substrate through which cue-related activation influence downstream striatal processing relevant to craving and relapse vulnerability (Goldstein et al., 2009; Goldstein & Volkow, 2011; Jentsch & Taylor, 1999). Consistent with this framework, early studies showed engagement of cingulate/ACC regions during exposure to drug-associated cues (Garavan et al., 2000; Zhang et al., 2011). Subsequent meta-analyses identified the ACC as a convergent node recruited across drug classes, together with striatal and insular regions (Jasinska et al., 2014; Kuhn & Gallinat, 2011; Noori et al., 2016). More recent meta-analyses further linked cue-induced activation in cingulate and insular regions to treatment outcomes (Evohr et al., 2025).

The ACC and broader frontostriatal circuitry are also engaged during attempts to regulate cue-induced craving. Cognitive down-regulation of craving is associated with increased recruitment of prefrontal control regions and reduced activity in craving-related nodes, including ventral striatum and cingulate/ACC-related circuitry (Kober et al., 2010). Multivariate analyses further identified distributed brain patterns linking cingulate/ACC and ventromedial prefrontal regions with subjective craving (Koban et al., 2023). While these human studies link ACC-centered circuits to cue-induced craving, preclinical models can causally manipulate homologous circuits and dissect their contributions to relapse-related behavior.

In the present study, we combined a human-relevant rat model of relapse after voluntary abstinence induced by adverse consequence of drug seeking with focal ACC-targeted TMS and rodent fMRI. We observed a significant reduction in the incubation of oxycodone seeking, underlying the behavioral relevance of ACC-centered circuits for relapse vulnerability in a clinically relevant abstinence context. We further examined how ACC-targeted TMS modulated frontostriatal functional connectivity during abstinence. In sham-treated rats, voluntary abstinence was associated with a reduction in ACC functional connectivity with both dorsal striatum and nucleus accumbens, major striatal targets of ACC projections within the corticostriatal circuits implicated in motivated behavior (Di Martino et al., 2008; Goldstein et al., 2009; Goldstein & Volkow, 2011; Haber & Knutson, 2010). In contrast, active TMS targeting the ACC prevented this time-dependent decrease in ACC-striatal connectivity. The incubation-related ACC-dorsal striatum connectivity changes are consistent with our prior findings that DMS inactivation reduced relapse after electric barrier-induced abstinence and increased ACC-DMS functional connectivity (Ma et al., 2025).

Together, our results showed that focal ACC neuromodulation can causally influence the corticostriatal network dynamics and relapse-like behavior in a clinically relevant animal model. By integrating circuit-resolved functional connectivity with targeted neuromodulation in rodents, our study extends correlational evidence from human neuroimaging and establishes a translational platform for linking network-level biomarkers of craving and relapse to mechanistically informed TMS interventions, with implications for target selection in clinical neuromodulation.

### TMS as a method for neuromodulation in rodent models and translational implications

Noninvasive brain stimulation is an established tool for modulating neural activity and connectivity in humans (Nitsche & Paulus, 2000; Pascual-Leone et al., 1996). Clinical studies have shown that repetitive TMS (rTMS) targeting the medial prefrontal cortex, ACC, or insular cortex reduces alcohol (Harel et al., 2022) or nicotine (Ibrahim et al., 2023) craving. However, the underlying mechanisms are unknown, and TMS has primarily been applied as an adjunctive rather than a stand-alone treatment (Tsai et al., 2021), with heterogenous outcomes reported across targets and stimulation protocols (Dinur-Klein et al., 2014; Hanlon et al., 2016; Luigjes et al., 2019; Steele & Maxwell, 2021). Thus, the development of a small-animal-compatible TMS system provides a critical opportunity to bridge preclinical and clinical research by enabling mechanistic examinations of stimulation effects.

In the present study, we addressed several key technical barriers that have previously limited the use of TMS in small animal models of addiction by using our custom-designed rat-specific TMS coil. First, our rat-specific TMS coil achieved a stimulation focality of approximately 2 mm, a level of focality comparable to that used in clinical applications (Meng et al., 2018). This enables more precise targeting of rodent brain regions with minimal off-target effects. Second, our hdTBS procedure enhances the stimulation aftereffects by ∼92% compared to conventional intermittent TBS (Meng et al., 2022). Third, we performed the stimulation in awake rats (Cermak et al., 2020), eliminating anesthesia-related confounds. Finally, we combined the novel TMS coil and hdTBS with rat fMRI to longitudinally track functional connectivity changes and assess TMS-induced effects, enabling investigation of mechanisms underlying the ‘treatment-like’ effects of TMS in the animal model (Ma et al., 2025). Collectively, this platform establishes a broad translational framework for dissecting neuromodulation mechanisms in rodent model of addiction and other psychiatric disorders, thereby informing network-guided target selection and protocol optimization for clinical TMS neuromodulation.

Our study further highlights a key translational advantage of using a focal TMS coil in rodent models by enabling direct integration of stimulation-induced functional changes with independently defined axonal projection architecture. By comparing TMS-induced functional connectivity changes with tracer-based axonal projection maps from ACC, we demonstrated that regions exhibiting significant stimulation-dependent interaction effects preferentially overlapped with regions receiving dense axonal projections from the stimulation site. This directional projection-level correspondence provides mechanistic insight into how focal stimulation propagates through a distributed network, an aspect that is not readily available in human studies. This integrative framework strengthens the interpretation of preclinical TMS findings and informs network-guided target selection for clinical neuromodulations. This integration is particularly valuable because diffusion MRI-based tractography, the primary approach available in humans, offers indirect, large-scale estimates of anatomical connectivity. In contrast, tracer-defined axonal projection provides complimentary directional information and shows closer correspondence with functional network organization (Melozzi et al., 2019; Schilling et al., 2019).

Consistent with the therapeutic potential of neuromodulation strategies, clinical studies have shown that high-frequency PFC rTMS reduces cue-induced opioid craving (Tsai et al., 2021) and cigarette consumption and craving (Steele et al., 2019). Deep TMS protocols targeting mPFC and ACC using the H-coil have shown reductions in cocaine (Martinez et al., 2018), alcohol (Harel et al., 2022), and nicotine (Dinur-Klein et al., 2014) intake, with the latter study additionally stimulating the insular cortex. These studies highlight the clinical relevance of ACC-related prefrontal networks as neuromodulation targets and provide clinical evidence for our reverse-translation framework that uses animal models mimicking effective human treatments to improve translational relevance and identify novel treatments (Venniro et al., 2020). A challenge for future research is to further advance coil design and stimulation procedures to improve the precision and depth of targeting deeper midline structures, thereby expanding their therapeutic potential in clinical settings (Deng et al., 2014; Nurmi et al., 2021).

### Methodological considerations and future studies

Our study has several limitations. First, we used only male rats. We made this choice because fMRI measures are sensitive to sex differences in brain anatomy and functional connectivity (Sumiyoshi et al., 2017; Wang et al., 2019), as well as ovarian hormone– and cycle-related fluctuations in functional connectivity (Filippi et al., 2013; Pritschet et al., 2020). In addition, our fMRI protocol requires light anesthesia (low-dose isoflurane combined with dexmedetomidine), for which sex- and estrous cycle– dependent responses have been reported (Vincent et al., 2023; Wasilczuk et al., 2024). Including females would therefore require different anesthetic dosing and data collection across estrous and non-estrous phases, substantially increasing variability and nearly tripling sample sizes. In this context, prior studies from our group reported no significant sex differences in oxycodone self-administration, electric barrier– induced suppression of drug intake, or incubation of oxycodone craving (Fredriksson et al., 2020; Fredriksson et al., 2023; Negishi et al., 2025). Nevertheless, we plan to examine the effects of hdTBS in female rats in future studies, as evidence suggests that even when relapse- and craving-related behavioral measures are similar across sexes, the underlying neural mechanisms may differ (Fredriksson, Venniro, et al., 2021; Nicolas et al., 2022).

Second, we did not evaluate the durability of the TMS treatment response. We exposed the rats to daily TMS during late abstinence and assessed drug seeking a day after the final stimulation session. Future studies should assess drug-seeking behavior at longer post-treatment intervals to determine the persistence of TMS effects.

Third, we derived axonal projection data from mouse anterograde tracer experiments, whereas we conducted fMRI and TMS modulation in rats. As such, the projection maps provide anatomical context but do not offer species-matched or causal validation of the observed functional connectivity changes.

Finally, we evaluated TMS effects using an animal model that captures a specific feature of human addiction: relapse after prolonged abstinence driven by re-exposure to drug-associated cues and contexts (Chow et al., 2025; O’Brien, 1997). An important direction for future research is to determine whether these findings generalize to other common relapse triggers in humans, such as stress (Sinha et al., 2011) and drug re-exposure (de Wit, 1996), which have also been extensively studied in animal models (Mantsch et al., 2016; Shaham et al., 2003).

## CONCLUSION

In summary, we applied focal TMS to the rat ACC using a hdTBS protocol in a model of relapse to opioid seeking. In sham-treated rats, incubation of oxycodone seeking was associated with a reduction in ACC functional connectivity, as revealed by resting-state fMRI. In contrast, daily hdTBS for seven days prevented both the incubation effect and the prefrontal connectivity alterations. TMS is a well-established, non-invasive neuromodulation technique with an excellent safety profile. Thus, our findings highlight a promising circuit-based strategy for relapse prevention.

## MATERIALS AND METHODS

### Subjects

We collected data from 36 adult male Sprague–Dawley rats (280–350 g before surgery) obtained from ENVIGO (Indianapolis). We initially group-housed the rats and individually housed them after surgery. We maintained the rats on a reverse 12-h:12-h light/dark cycle with ad libitum access to food and water. The Animal Care and Use Committee of the National Institute on Drug Abuse Intramural Research Program approved all experimental procedures.

We previously found no sex differences in oxycodone self-administration, electric barrier–induced suppression of drug intake, or incubation of oxycodone craving (Fredriksson et al., 2020; Fredriksson et al., 2023; Negishi et al., 2025). Therefore, we used only male rats to minimize variance in fMRI-based measurements of brain functional connectivity, which has been documented to show sex differences (Sumiyoshi et al., 2017; Wang et al., 2019), and ovarian hormone–dependent, cycle-related fluctuations (Filippi et al., 2013; Pritschet et al., 2020). Inclusion of females rats would introduce additional data variability related to estrous cycle-dependent changes in functional connectivity and would require a substantial increase in sample size and cycle tracking to achieve comparable statistic power. Given the scope and technical demands of the current longitudinal TMS-fMRI design, including both sexes was not feasible.

### Drugs

We obtained oxycodone hydrochloride (HCl) from the NIDA pharmacy and diluted it in sterile saline at a unit dose of 0.1 mg/kg for self-administration training. We selected this unit dose based on our previous studies (Fredriksson et al., 2020; Fredriksson et al., 2023; Ma et al., 2025; Negishi et al., 2025).

### Surgery

As described previously (Ma et al., 2025), we implanted customized intravenous catheters into the jugular vein under isoflurane anesthesia (3–4% induction, 1.5–2.5% maintenance). After intravenous surgery, we transferred the rats to a stereotaxic frame (Kopf Instruments) and head-fixed them for transcranial headpost implantation. We carefully implanted a 3D-printed, T-shaped headpost onto the rat skull to serve as a reference for accurate coil positioning (Cermak et al., 2020). We implanted two to three PEEK screws (NBK America LLC) and applied dental cement (C&B Metabond, Parkell) to the skull surface to secure the headpost. When necessary, we added a detachable 3D-printed coil guide of variable thickness to further adjust coil positioning. We administered ketoprofen (2.5 mg/kg, subcutaneous) twice, at surgery and 24 h later, for analgesia and anti-inflammatory treatment. We flushed the intravenous catheters with gentamicin (4.25 mg/ml) and heparin (30 unit/ml) every 24–48 h until the end of the experiments.

### Behavioral experiments

#### Oxycodone self-administration

After 7–10 days of recovery from surgery, we trained the rats to self-administer oxycodone on a fixed-ratio 1 (FR1) schedule with a 20-s timeout for 6 h/day over 14 days (Figure 1A). Each session began with illumination of a red houselight followed by insertion of the active lever. Active lever presses delivered one oxycodone infusion (100 µl over 3.5 s; 0.1 mg/kg). Each infusion was paired with a 20-s discrete tone and illumination of a white cue light above the active lever. During the 20-s timeout, active lever presses were recorded but not reinforced. Inactive lever presses had no programmed consequences but were recorded. We limited the maximum number of infusions to 90 per 6-h session. At the end of each session, the active lever was retracted, and the houselight was turned off.

#### Electric barrier–induced abstinence

Rats underwent electric barrier-induced voluntary abstinence for 13 days (Figure 1A). During this phase, oxycodone was available for 2-h/day under the same dose, reinforcement schedule, and tone–light cue conditions used during self-administration training. We induced voluntary abstinence by introducing an electric barrier (“shock zone”) that covered two-thirds of the chamber floor near the drug-paired active lever (Fredriksson et al., 2020; Ma et al., 2025). When rats approached the drug-paired lever, they received a continuous mild footshock through the grid floor. We increased shock intensity daily from 0.0 mA to 0.3 mA in 0.1 mA increments. For rats that did not suppress oxycodone self-administration (<3 infusions/session), we increased the shock intensity up to 0.4 mA.

#### Relapse test

We assessed relapse to oxycodone seeking on day 1 (30-min test) and day 15 (90-min test) under extinction conditions (Figure 1A). We turned off the electric barrier on relapse test days. On testing days, we habituated the rats to the chamber for 30 min before session onset to allow exploration. During relapse tests, active lever presses activated the infusion pump and the drug-paired tone-light cue but did not deliver oxycodone.

#### Data analysis

We analyzed behavioral data using repeated-measures ANOVAs in SPSS 29. When main effects or interactions reached significance (p < 0.05), we conducted post-hoc comparisons using Fisher’s LSD. We describe the specific between- and within-subject factors in the Results section. For clarity, we report only statistical effects critical for data interpretation and indicate significant post-hoc comparisons with asterisks in the figures (see Table S1 for a complete summary of statistical analyses).

### TMS experiments

We used a custom designed and built TMS coil with an approximate focality of 2 mm (Figure 1D–E) (Meng et al., 2018). Because this high focality requires precise targeting, we implanted a headpost on the rats’ skull to serve as a reference for accurate and consistent coil placement during TMS delivery.

#### Habituation

Before TMS delivery, we habituated the rats for 6 days to the TMS coil to minimize stress and movement. We handled the rats, held them awake under the TMS coil, and played recorded hdTBS sounds (Figure 1C). We repeated this procedure three times per day. After habituation, the rats showed reduced acute stress responses including decreased vocalization, urination, defecation, and escape attempts.

#### TMS delivery

After habituation, we assessed motor threshold in each rat before TMS treatment. We defined motor threshold as the minimum TMS power that evoked hindlimb motor responses in 5 of 10 pulses delivered to the corresponding motor cortex (Cermak et al., 2020; Meng et al., 2022). We used the motor threshold to set individual TMS power levels. In the hdTBS group, we set TMS power to 125% of motor threshold and positioned the rat’s head close to the coil surface (Figure 1C). In the sham group, we set TMS power to 75% of motor threshold and inserted a 4-cm plastic spacer between the rat head and coil surface (Figure 1C). This configuration reduced the electric field near the skull to near zero while maintaining comparable acoustic stimulation. We manually blocked the rats’ ears to minimize the effects of the acoustic noise from the TMS coil. Each session consisted of 6 pulses delivered at 50 Hz (20 ms interpulse interval), with 10 bursts per train delivered at 5 Hz (200 ms interbursts interval). We delivered 20 trains (2 s each) separated by 8 s intertrain intervals, resulting in 1200 total pulses over 192 s per session. We applied dry ice to prevent coil overheating and allowed a 60-min cooling period between rats to ensure complete coil cooling.

### fMRI experiments

#### Animal preparation

We anesthetized rats for fMRI using a combination of isoflurane and dexmedetomidine as described previously (Brynildsen et al., 2017; Lu et al., 2012). We initially anesthetized the rats with 2.5% isoflurane in oxygen-enriched air (70% N_2_, 30% O_2_), followed by an intraperitoneal bolus of dexmedetomidine (0.015 mg/kg). We then transferred the rats to a customized MRI-compatible holder and head-fixed them. We gradually reduced isoflurane to 0.5–0.75% and delivered it through a nose cone. We delivered continuous subcutaneous dexmedetomidine infusion (0.015 mg/kg/h) using an infusion pump (PHD 2000, Harvard Apparatus). We monitored physiological parameters throughout scanning and initiated fMRI data acquisition at least 60 min after anesthesia induction (Brynildsen et al., 2017). We maintained body temperature with a temperature-controlled water-heating pad and monitored blood oxygenation and respiration (SA Instruments). We adjusted oxygen concentration to maintain SpO_2_ ≥ 95%.

#### Data acquisition and analysis

We acquired MRI data using a Bruker Biospin 9.4T scanner with Paravision 7.0 software. We used a volume quadrature transmitter coil (MT0381) for radiofrequency excitation and a circular surface coil (MT0105-20) for signal reception. We used the decussation of the anterior commissure (∼ −0.36 mm from bregma) as a fiducial landmark to standardize slice localization across rats (Lu et al., 2007). We collected high-resolution structural images using a RARE sequence (TR = 3100 ms, TE = 36 ms, FOV = 30 × 30 mm^2^, matrix = 256 × 256, slice thickness = 0.6 mm, gap = 0.1 mm, 31 slices). We acquired resting-state fMRI data using an in-house gradient-echo EPI sequence with forward and reverse k-space trajectories. We corrected geometric distortions in EPI images using the reverse k-space trajectory methods (Chang & Fitzpatrick, 1992). Scan parameters were TR = 1500 ms, TE = 15 ms, FOV = 30 × 30 mm^2^, matrix = 80 × 80, slice thickness = 0.6 mm, gap = 0.1 mm, and 19 slices.

We processed resting-state fMRI data using established pipelines (Lu et al., 2012), including distortion correction, skull stripping, registration to a standard template, independent component analysis (ICA)-based noise removal, band-pass filtering (0.01–0.1 Hz), and spatial smoothing (full width at half maximum = 0.8 mm) using Analysis of Functional NeuroImages (AFNI, https://afni.nimh.nih.gov/) (Cox, 1996), Advanced Normalization Tools (ANTs, http://stnava.github.io/ANTs/) (Avants et al., 2011), and FMRIB Software Library (FSL, https://fsl.fmrib.ox.ac.uk) (Smith et al., 2004).

We conducted seed-based functional connectivity analyses by extracting and averaging voxel time courses from predefined seed regions and correlating them with whole-brain time courses. We converted correlation coefficients to z-scores. We then performed voxelwise Group (sham, hdTBS) × Abstinence Time (early, late) mixed-effects analyses using AFNI’s 3dLME, followed by post-hoc t-tests. We applied multiple-comparison correction using 3dClustSim (Cox et al., 2017). We considered clusters significant at uncorrected p < 0.01 with a minimum cluster size of 25 voxels. We mapped significant clusters to anatomical regions using a rat brain atlas (Paxinos, 2007) and extracted voxel-wise connectivity values from each ROI for further analysis.

### Axonal projection analysis

We obtained tracer-based axonal projection data from an anterograde viral tracer experiment involving ACC AAV-EGFP injections from the Allen Mouse Brain Connectivity Atlas (Oh et al., 2014) (Experiment ID: 112458114). We downloaded voxel-wise projection images at 25 µm isotropic resolution and regional projection summary metrics in the Allen Common Coordinate Framework (CCFv3). Since stereotaxic coordinates are not directly interchangeable between rat and mouse brains, we did not match the anterior to posterior (AP) levels based solely on bregma coordinates. Instead, we aligned rat fMRI slices to the Allen mouse atlas using conserved anatomical landmarks, including the morphology and relative position of the corpus callosum, anterior commissure, lateral and third ventricles, and striatal boundaries. After landmark-based alignment, we used the corresponding Allen z-index to compute the approximate AP position relative to an empirically determined bregma location. Bregma in the 25 µm CCF was approximate at z = 218, and the AP position was calculated as:

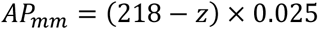

We excluded regions with negligible or zero projection values from further analysis. For quantitative analyses, we extracted two regional projection measures:

Projection volume = Volume of projection signal in structure in mm^3^.

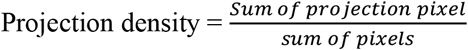

## List of Supplementary Materials

Figure S1 to S2.

Table S1.

## Acknowledgments

This work was supported by NIDA-NIH Intramural Research Program funds.

## Author contributions

Conceptualization: ZM, YD, ES, ZX, YS, HL, YY

Methodology: ZM, YD, HN, SL, MH, YS, HL, YY

Investigation: ZM, YD, SL, SH, AC, TS, OV, HL

Data analysis: ZM, YD, SL, DW

Visualization: ZM, YS, YY

Writing – original draft: ZM, YS, HL, YY

Writing – review & editing: All authors

## Data and materials availability

All data and materials associated with this study are present in the paper or the Supplementary Materials.

