## Supplementary Materials for "Focal Transcranial Magnetic Stimulation of the Rat Anterior Cingulate Cortex Inhibits Incubation of Opioid Craving after Voluntary Abstinence"

**Supplementary figures:**

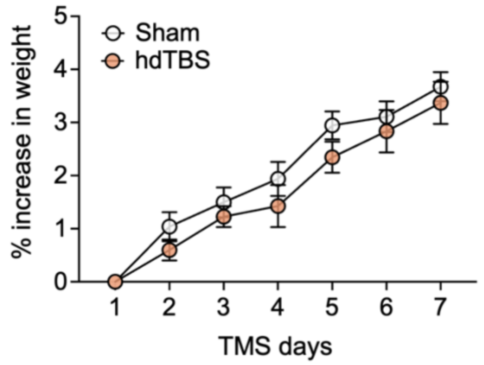


**Figure S1. Percentage of weight increase during TMS.** We analyzed the percentage of body weight increase during the 7 TMS days with RM-ANOVA using between-subject factor of Group (Sham, hdTBS) and within-subject factor of TMS days (day 1-7). Normal increase in body weights was observed in both sham and hdTBS rats as indicated by significant effect of TMS days ($F_{6, 204} = 78.04$, $p < 0.001$, Table S1). Data show mean ± SEM. n = 17 sham, n = 19 hdTBS.

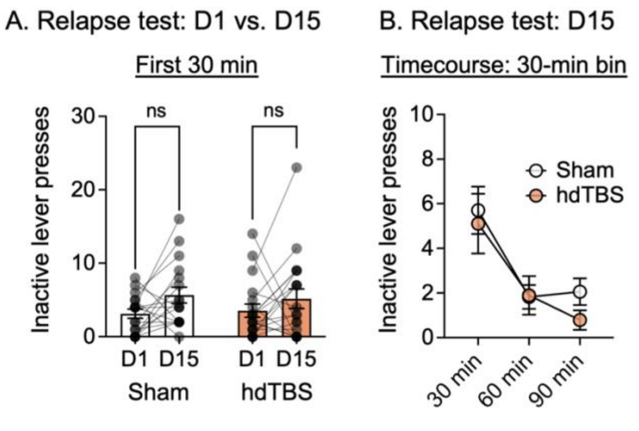


**Figure S2. Inactive lever presses during relapse test.** (**A**) Relapse test: day 1 vs. day 15. Data show mean ± SEM number of inactive lever presses during the 30-min test (day 1) and the first 30-min of test (day 15). (**B**) Relapse test day 15: 30 min bin timecourse of inactive lever press. Data show mean ± SEM. n = 17 sham, n = 19 hdTBS. See Table S1 for complete statistics.

### Supplementary tables

**Table S1. Statistical analysis for Figure 1.** Statistical analysis for the behavioral results (SPSS GLM repeated-measures module, GLM univariate module, and independent-samples T-test). Partial Eta[2] ($\eta_p^2$) = proportion of explained variance. Cohen's d ($d$) = standardized effect size. Observed power: computed using alpha = 0.05. RM: repeated measures.

| Figure number | F- or T-value | p-value | Effect size | Observed power |
|---|---|---|---|---|
| Figure 1B. Self-administration infusions<br>RM-ANOVA<br>Within-subject factor: Training Session<br>Between-subject factor: Group | Training Sessions (1-14) within-subject: $F_{13, 442} = 89.62$<br>Group (Sham, hdTBS) between-subject: $F_{1, 34} = 0.47$<br>Group × Training Session interaction: $F_{13, 442} = 1.47$ | < 0.001*<br>0.59<br>0.13 | $\eta_p^2 = 0.74$<br>$\eta_p^2 = 0.014$<br>$\eta_p^2 = 0.041$ | 1.00<br>0.10<br>0.82 |
| Figure 1B. Electric barrier infusions<br>RM-ANOVA<br>Within-subject factors: Electric Barrier Session<br>Between-subject factor: Group | Electric Barrier Sessions (1-13) within-subject: $F_{12, 408} = 104.25$<br>Group (Sham, hdTBS) between-subject: $F_{1, 34} = 1.63$<br>Group × Electric Barrier Session interaction: $F_{12, 408} = 1.40$ | < 0.001*<br>0.21<br>0.16 | $\eta_p^2 = 0.75$<br>$\eta_p^2 = 0.046$<br>$\eta_p^2 = 0.039$ | 1.00<br>0.24<br>0.77 |
| Figure 1F. Motor thresholds machine output<br>Two-sample T-test | Sham vs. hdTBS: t = 1.69, df = 34 | 0.10 | $d = 0.56$ | 0.37 |
| Figure 1H. Relapse test active lever press<br>RM-ANOVA<br>Within-subject factor: Abstinence Day<br>Between-subject factor: Group | Abstinence Day (Day 1, Day 15) within-subject: $F_{1, 34} = 13.72$<br>Group (Sham, hdTBS) between-subject: $F_{1, 34} = 5.62$<br>Group × Abstinence Day interaction: $F_{1, 34} = 4.40$ | < 0.001*<br>0.024*<br>0.043* | $\eta_p^2 = 0.29$<br>$\eta_p^2 = 0.14$<br>$\eta_p^2 = 0.12$ | 0.95<br>0.63<br>0.53 |
| Figure 1I. Relapse test (30 min bin timecouse) active lever press<br>RM-ANOVA<br>Within-subject factor: Session Time<br>Between-subject factor: Group | Sessions Time (30, 60, 90 min) within-subject: $F_{2, 68} = 35.83$<br>Group (Sham, hdTBS) between-subject: $F_{1, 34} = 7.36$<br>Group × Session Time interaction: $F_{2, 68} = 2.42$ | < 0.001*<br>0.010*<br>0.097 | $\eta_p^2 = 0.51$<br>$\eta_p^2 = 0.18$<br>$\eta_p^2 = 0.066$ | 1.00<br>0.75<br>0.47 |
| Figure S1. Body weight percentage change<br>RM-ANOVA<br>Within-subject factor: TMS Day<br>Between-subject factor: Group | TMS Day (1-7) within-subject: $F_{6, 204} = 78.04$<br>Group (Sham, hdTBS) between-subject: $F_{1, 34} = 1.24$<br>Group × TMS Day interaction: $F_{6, 204} = 0.48$ | < 0.001*<br>0.27<br>0.82 | $\eta_p^2 = 0.70$<br>$\eta_p^2 = 0.035$<br>$\eta_p^2 = 0.014$ | 1.00<br>0.19<br>0.19 |
| Figure S2. Relapse test inactive lever press<br>RM-ANOVA<br>Within-subject factor: Abstinence Day<br>Between-subject factor: Group | Abstinence Day (Day 1, Day 15) within-subject: $F_{1, 34} = 4.57$<br>Group (Sham, hdTBS) between-subject: $F_{1, 34} = 0.001$<br>Group × Abstinence Day interaction: $F_{1, 34} = 0.21$ | 0.040*<br>0.97<br>0.65 | $\eta_p^2 = 0.12$<br>$\eta_p^2 = 0.000$<br>$\eta_p^2 = 0.006$ | 0.55<br>0.050<br>0.073 |
| Figure S2. Relapse test (30 min bin timecouse) inactive lever press<br>RM-ANOVA<br>Within-subject factor: Session Time<br>Between-subject factor: Group | Sessions Time (30, 60, 90 min) within-subject: $F_{2, 68} = 13.72$<br>Group (Sham, hdTBS) between-subject: $F_{1, 34} = 0.59$<br>Group × Session Time interaction: $F_{2, 68} = 0.32$ | < 0.001*<br>0.45<br>0.73 | $\eta_p^2 = 0.29$<br>$\eta_p^2 = 0.017$<br>$\eta_p^2 = 0.009$ | 1.00<br>0.12<br>0.100 |
